# Systems profiling reveals recurrently dysregulated cytokine signaling responses in ER+ breast cancer patients’ blood

**DOI:** 10.1101/2023.10.31.564987

**Authors:** Brian Orcutt-Jahns, Joao Rodrigues Lima Junior, Russell C. Rockne, Adina Matache, Sergio Branciamore, Ethan Hung, Andrei S. Rodin, Peter P. Lee, Aaron S. Meyer

## Abstract

Cytokines mediate cell-to-cell communication across the immune system and therefore are critical to immunosurveillance in cancer and other diseases. Several cytokines show dysregulated abundance or signaling responses in breast cancer, associated with the disease and differences in survival and progression. Cytokines operate in a coordinated manner to affect immune surveillance and regulate one another, necessitating a systems approach for a complete picture of this dysregulation. Here, we profiled cytokine signaling responses of peripheral immune cells from breast cancer patients as compared to healthy controls in a multidimensional manner across ligands, cell populations, and responsive pathways. We find alterations in cytokine responsiveness across pathways and cell types that are best defined by integrated signatures across dimensions. Alterations in the abundance of a cytokine’s cognate receptor do not explain differences in responsiveness. Rather, alterations in baseline signaling and receptor abundance suggesting immune cell reprogramming are associated with altered responses. These integrated features suggest a global reprogramming of immune cell communication in breast cancer.

**Significance Statement:** While individual cytokine responses have previously been observed to be altered in breast cancer, cytokine signaling responses are tightly interconnected in a way that has not been previously characterized. Here, we profile cytokine signaling responses and find alterations that are shared across both pathways and cell types. The signatures across these measurements better define the alterations and point to a broad immunosuppression response.

**Highlights:**

- Baseline and post-stimulation cytokine signaling profiles differ between healthy donors and breast cancer patients.
- Changes in cytokine response are not explained by differences in abundance of the cognate receptor
- Features of signaling response and receptor abundance dysregulation are coordinated across patients
- Integrated patterns of dysregulation in breast cancer patients share features of Th17 like-response as well as regulatory-like B and CD8^+^ T cells

## INTRODUCTION

Cytokines are extracellular proteins that mediate cell-to-cell communication within the immune system. Many groups have observed altered levels of various cytokines in cancer^1^. In addition, we have observed that immune cell cytokine responsiveness is altered in cancer patients, leading to perturbed immune cell function and differentiation^2–5^. In estrogen receptor-positive breast cancer (ER+ BC, of any stage) patients, we found that ∼40% harbor defects in immune signaling at time of diagnosis, including altered signaling responses to IL-6, IFN-γ, TGF-β, IL-10, and IL-4 in various immune cells^6,7^. Importantly, these peripheral immune defects also reflect the tumor immune microenvironment and predict clinical outcome. Mechanisms that connect altered signaling responses to dysfunctional antitumor immunity remain poorly understood^6,8^. If a mechanistic understanding of these defects can be realized, therapeutically correcting these defects may offer a novel approach for improving outcomes in ER+ BC patients.

Because cytokines operate in combination in vivo, defining their integrated pathway responses and downstream consequences is a major unresolved issue in immune signaling and cancer biology. Thus, there is a major gap in our understanding of how the immune system functions from the perspective of cytokine-mediated information flow, and in our ability to predict how alterations might hinder or drive its disruption. Immune signaling defects in BC patients can be observed through perturbation experiments, wherein patients’ peripheral blood mononuclear cells (PBMCs) are treated with different cytokines, and their signaling responses are measured^2,3,6,7^. These rich data reveal abnormalities in signaling responses which can be reflected in the dynamics of cytokine response for one or several cytokines and cell populations, with heterogeneity across subjects. Thus, while these data are sufficiently rich to uncover defects in immune signaling, the richness itself presents challenges. A central challenge is that cytokines responses vary across several dimensions—depending on the cytokine ligand, temporal dynamics, cell type, patient, and responsive signaling pathway. Exploring these in combination creates an explosion of variables through their pairwise combination, and typical dimensionality reduction methods (e.g., PCA, t-SNE, U-MAP) are have limitations in exploring the data^9^.

Dimensionality reduction in tensor form, wherein data is organized into a multi-dimensional array, can preserve the natural organization of profiling experiments that thoroughly characterize these dimensions of cytokine response^9^. When data can be arranged in tensor form, tensor decomposition methods can provide improvements in interpretation of the resulting factors, reduce the data to a greater extent, handle missing values and batch effects, and improve the ability to draw associations (e.g., between patterns and BC status)^9–11^. These properties arise from the ability of tensor decompositions to define the association of each component pattern with each dimension separately^10–13^. Consequently, these methods avoid repetition of pattern effects over variables that measure shared features—for instance, in measuring temporal patterns, various signals can be defined as a single temporal signature^13^.

By defining component associations with respect to each dimension, tensor decomposition methods are especially effective in integrating datasets of diverse structures with one or more shared dimensions, allowing common patterns to be identified^10,14–17^. As linear methods, tensor decompositions have well-defined properties of convergence, solving, scalability, and interpretation, like matrix-based approaches such as principal component analysis (PCA) or non-negative matrix factorization (NNMF) ^9^. However, unlike matrix-based decompositions, tensor decompositions exist in a variety of structural forms. Other approaches such as generalized SVD, higher-order SVD, and others have comparative benefits in the compactness of the resulting decomposition or numerical properties, sometimes at the cost of easy interpretation^18^. Development of new algorithms, decomposition structures, and interpretation/visualization approaches, and accessible software implementations are all areas of very active research^12,19,20^.

Here, we profile the cytokine responses of immune cells in the peripheral blood from breast cancer patients as compared to healthy controls in a multidimensional manner across ligands, cell populations, and responsive pathways via high-dimensional (18- and 24-color) phosflow analysis. This allows us to quantify major pSTATs concurrently within individual immune cell subsets. We find alterations in cytokine responsiveness across pathways and cell types are widespread in BC compared to healthy subjects, and these changes can be described by signatures across cell types, pathways, and cytokines. Tensor decomposition provides a way to define these signatures across these dimensions and allows us to define these changes in an integrative manner. With this more complete view, we observe that alterations in the breast cancer cohort’s baseline signaling, receptor amounts, and cytokine responsiveness across cell types exhibit several coordinated features of dysregulated immune signaling, such as the simultaneous presence of a Th17 like-response, or B and CD8^+^ T cells displaying regulatory-like phenotypes.

## RESULTS

### Systematically identifying immune signaling response patterns

To systematically characterize the patterns of immune signaling response and dysregulation in ER^+^ breast cancer patients, we analyzed peripheral blood mononuclear cells (PBMCs) from 36 subjects, 22 of which were healthy donors, and 14 of which were newly diagnosed with ER+ breast cancer (Fig. 1a). We systematically treated the PBMCs with 7 different unique cytokine treatments to profile how the two cohorts differ in their signaling response. The cytokines included IL-2, IL-4, IL-6, IL-10, IFNγ, and TGFβ, as well as a combination of IFNγ and IL-6. After treatment for 15 minutes, cells were fixed and stained for 27 different intra- and extracellular markers and then gated into 23 distinct populations. Alongside various cell type markers, the phosphorylation responses of key transcription factors were quantified (pSTAT1, pSTAT3, pSTAT4, pSTAT5, pSTAT6, and pSmad2/3).

**Fig. 1.**
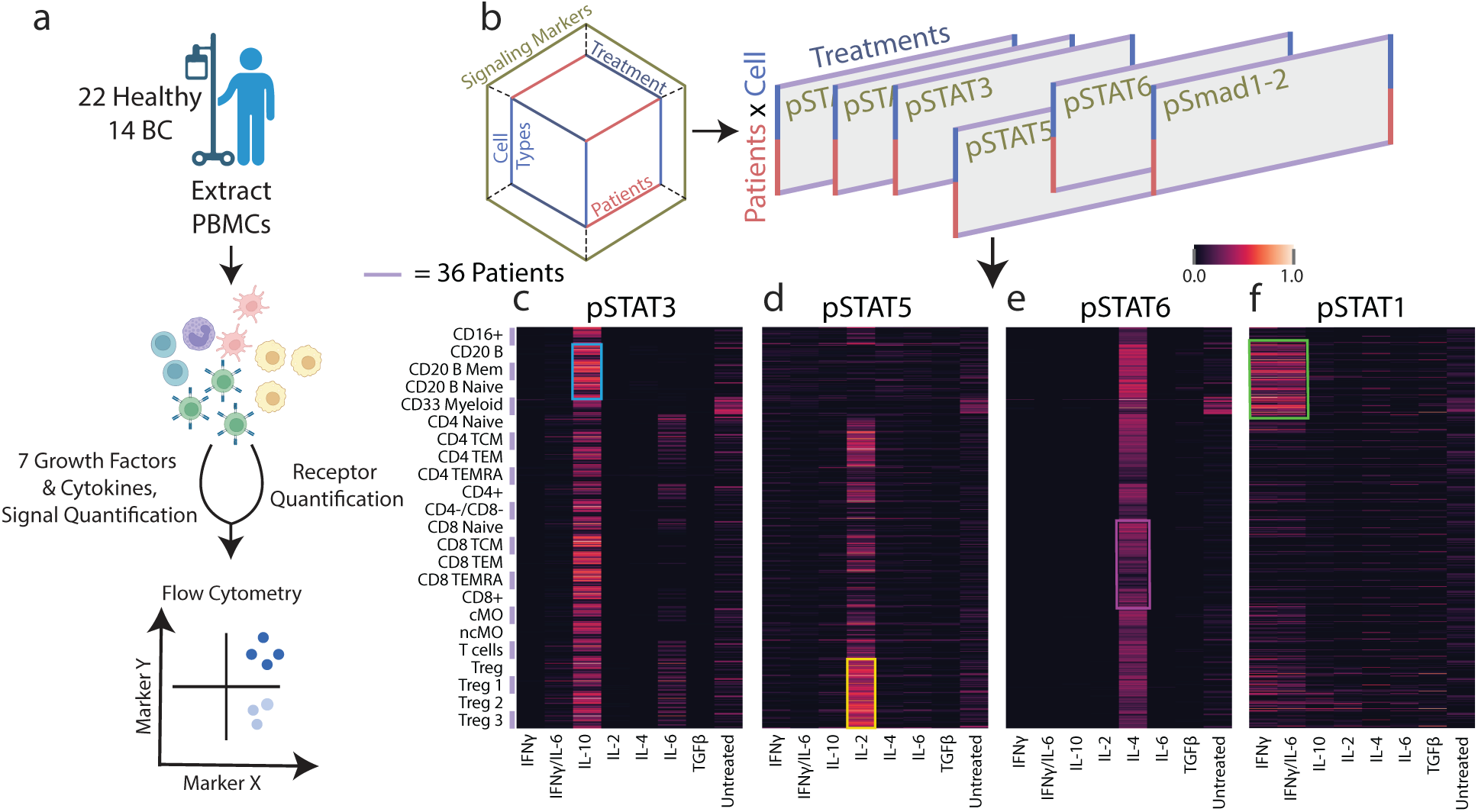
Systematically profiling PBMC cytokine signaling across several dimensions. **(a)** Schematic of the experimental approach. Human PBMCs were harvested from healthy and BC cohorts and subsequently treated with a panel of 7 cytokine/growth factor combinations. Response was quantified using 27-channel flow cytometry, through which the response of 23 different cell types was quantified. **(b)** Schematic of structure of the dataset and how it was divided for visualization in c–f. **(c– f)** Heatmap of phosphorylated STAT3 (c), STAT5 (d), STAT6 (e), and STAT1 (f) measurements for each treatment (X axis), and patient/cell type pair (Y axis). The signal was normalized to the maximum observed signal across patients. A small number of missing values were imputed for select patients in response to TGFβ and the IFNγ/IL-6 combination.

Using these data, we first confirmed that several canonical signaling trends were captured. We isolated subsets of our data and collapsed the data into matrices to visualize how cells from different patients (Y axis) responded to each cytokine treatment (X axis) as measured by STAT3, STAT5, and STAT6 phosphorylation (Fig. 1b–e). For instance, IL-10 stimulation resulted in STAT3 phosphorylation, as expected, that was most abundant in B cells (Fig. 1c, blue box)^21^. IL-2 induced STAT5 phosphorylation and was found most abundantly across regulatory T cells (T_reg_s), which are known to be sensitive to IL-2 stimulation (Fig. 1d, yellow box). STAT6 was found to be most strongly phosphorylated by IL-6; however, there was substantial variation in response within CD8^+^ cells across patients (Fig. 1e, purple box). Finally, STAT1 phosphorylation was especially responsive to IFNγ stimulation in B cells and CD33-positive myeloid cells (Fig. 1f, green box). However, to integrate each pattern of response and their association with time points, treatments, cell types, signaling pathways, and patients, a systems-level approach to the analysis was required.

### CPD reveals coordinated and substantial BC-associated patterns of signaling response

Examining individual cytokine responses or responsive pathways can only provide a limited picture of how responses vary, which is a common challenge with profiling data collected across several experimental variables/dimensions^22^. Therefore, we sought to identify patterns of response variation using an approach that explicitly accounts for data gathered across multiple experimental conditions. We applied canonical polyadic decomposition (CPD), a tensor decomposition method which allowed us to visualize the variation across each patient, treatment, cell type, and signaling pathway, or dimension. CPD factors an n-dimensional tensor, in this case four-dimensional, into the sum of vector outer products (Fig. 2a). Analogous to matrix-based methods like PCA, each set of vectors represents a pattern/component, and vectors associate with a specific dimension, describing how the pattern is represented along that axis. As a result, each factor describes how patterns vary along a specific dimension, and each component encodes a distinct response pattern. We applied CPD to our signaling data, which allowed us to preserve the patient, cell type, treatment, and signaling marker dimensions by avoiding the need to flatten the dataset into two dimensions, as needed for 2D matrix factorization methods like NMF or PCA. About 80% of the variance in our original dataset could be captured with 12 components (Fig. 2b). CPD led to a more concise representation of the data compared to PCA: 12 components were still more concise than a 1-component PCA model and explained a similar amount of the data compared to PCA at less than 1% of the size (Fig. 2c). We confirmed the appropriateness of CPD as a representation of the data and that the size of the factorization is appropriate by evaluating the imputation capability of the model when leaving out random values or patient-treatment-marker chords of the data. CPD was shown to accurately impute missing data up to and past 12 components, showing that the CP decomposition was able to recover and recapitulate missing data effectively, and that there were no superfluous or redundant components at such decomposition ranks (Fig. S2).

**Fig. 2.**
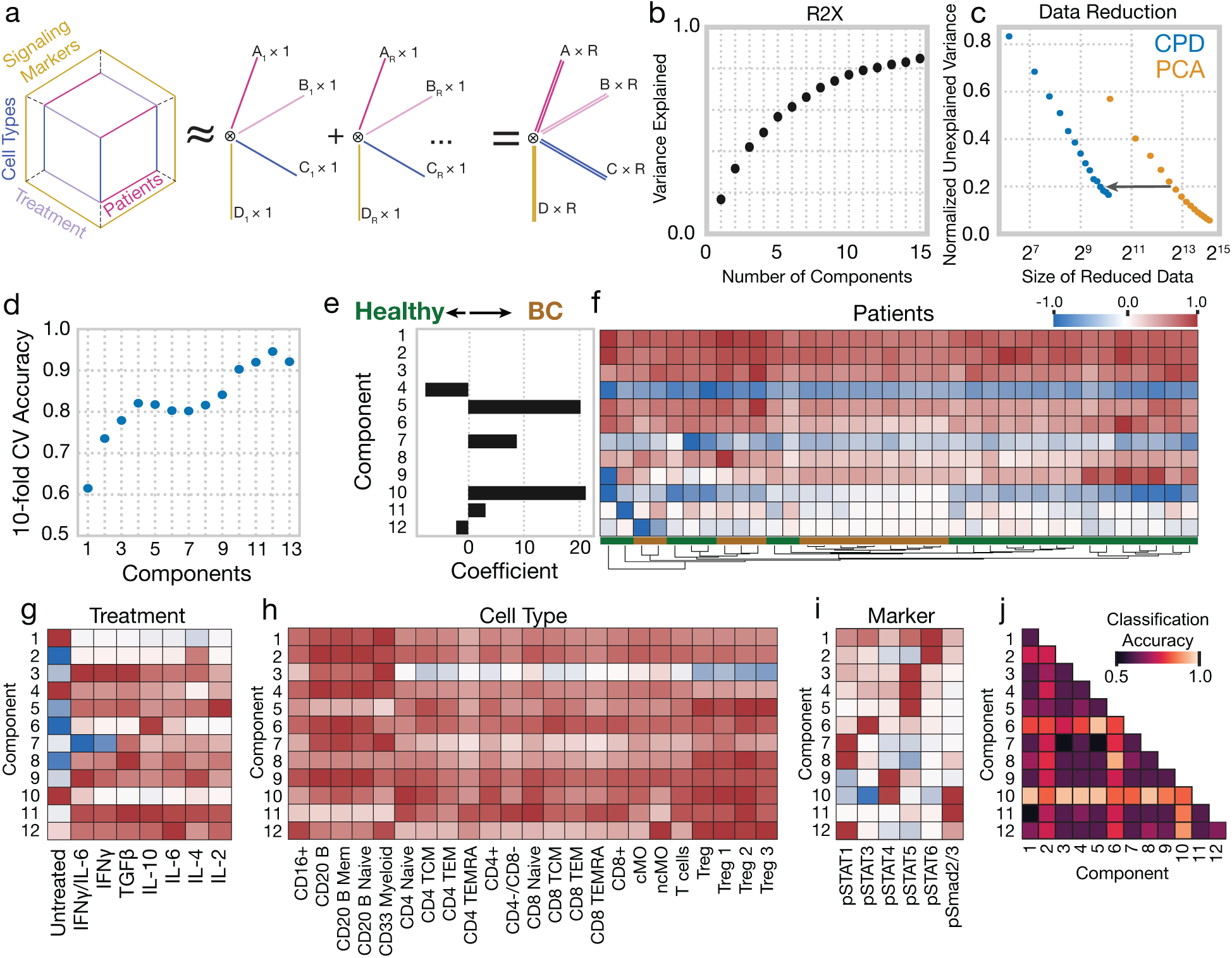
Canonical polyadic decomposition (CPD) of cytokine response identifies several response patterns strongly associated with BC. **(a)** Schematic of the CPD. The signaling data is organized into a four-dimensional tensor with axes for each patient, treatment, cell type, and signaling marker. This tensor is reduced into the sum of the outer products of vectors (components) associated with each dimension. **(b)** Percent variance reconstructed (R2X) versus the number of components used. **(c)** The remaining error on reconstruction, normalized to the total dataset variance, versus the size of the dataset after decomposition using CPD or PCA. **(d)** The accuracy of a logistic regression classifier upon 10-fold cross-validation, using the CPD patient factors with varying numbers of components. **(e)** The weights associated with each component for a logistic regression classifier fit to patient factors using a 12-component decomposition (healthy = 0, BC = 1). **(f)** A heatmap of the patient factor matrix, with the patients hierarchically clustered. Patient status is indicated by the coloring along the bottom. **(g–i)** Component values for each treatment (g), cell type (h), and signaling marker (i). **(j)** Classification accuracy for logistic regression classifiers using all pairs of components.

While the factors for each dimension coordinately explain the data, the patient factors specifically describe how these patterns vary across the patients. As the number of components is a tunable setting within CPD to explain more or less of the variation, we identified which number of components best predicted disease status to select an appropriate rank for our decomposition (Fig. 2d). With logistic regression, 12 components were optimal, with 94% accuracy on cross-validation (Fig. 2d). A similar modeling approach using fold-changes in signaling response was similarly predictive (Fig. S3). Interestingly, when using a similar approach with cell type abundances as inputs, we were unable to identify patterns associated with BC status, suggesting that no significant differences in the composition of cell types exist between the healthy and BC cohorts (Fig. S4). Next, we examined how the CPD-defined patterns associated with patient status through the logistic regression weights. Components 2, 5, 9, and 10 were most informative of patient disease status (Fig. 2e). Examining the patient factors overall, we found that that clusters of patients formed largely according to disease status; however, multiple clusters of both healthy and diseased patients were uncovered, suggesting that variation even within these cohorts was present (Fig. 2f).

The CPD factors effectively mapped expected signaling trends to specific patients, treatments, cell types, and signaling markers (Fig. 2f–i). One can readily interpret the resulting factor plots by tracing an individual component across each dimension. For instance, component 2 specifically associates with IL-4 treatment (Fig. 2g) and STAT6 phosphorylation (Fig. 2i). This pattern occurs in all patients (Fig. 2f) and all cell types, though more so in B cells (Fig. 2h). Furthermore, greater IL-4-mediated stimulation of pSTAT6 is associated with a lower baseline level of the marker (Fig. 2g). Thus, we can infer that component 2 summarizes STAT6 phosphorylation by IL-4, across patients and many cell types, but particularly in B cells. Another canonical response summarized by CPD is component 7. This component is more variable among patients (Fig. 2f), occurs in response to IFNγ (Fig. 2g), principally in B and myeloid cells (Fig. 2h), and represents the phosphorylation of STAT1 (Fig. 2i). While our regularized multivariate logistic regression classifier selected these components, we further examined whether components which may be predictive of disease status may be regularized out during the fitting process. By examining the classification accuracy of using pairwise combinations of components, we additionally found that component 6, which represented STAT3 phosphorylation in response to IL-10, was among the components which separated BC patients from healthy patients the best (Fig. 2j). Furthermore, this accuracy was much enhanced when component 6 was considered in combination with component 5.

Taken together, CPD effectively reduced the response data into coordinated patterns of cytokine response, which in turn associated strongly with BC status. CPD identified that BC patients are most prominently distinguished by their basal and induced pSmad2/3 and pSTAT4 across many populations (component 10), reduced IL-10 response across many cell populations (component 6), heightened IL-2 response in several populations but most prominently regulatory T cells (component 5), and to a lesser extent increased IFNγ response, mostly in B and CD33+ myeloid cells (component 7) (Figs. 2f–j, S5).

### Immune dysregulation is more prominently reflected in patterns of signaling response changes

After finding that patient-associated factors were overall predictive of disease status, we sought to dissect the specific changes associated with BC. While we could observe the prominent difference in IL-10-induced pSTAT3 in a per-measurement analysis (Fig. 3a), we wondered whether the CPD results offer additional insight into this prominent difference. First, we noted that the component associated with this reduction in response was also associated with an increase in baseline pSTAT3, despite only a few cell types featuring statistically significant difference in baseline pSTAT3 associated with BC (Fig. 3b). However, comparing the baseline versus induced pSTAT3 in CD8 cells, we indeed observed that, across patients, increased baseline pSTAT3 was associated with reduced response (Fig. 3c). Second, each component should represent coordinated patterns that are shared across cell types and patients (Fig. 2g). To examine this, we plotted the correlation in response between B cells and CD8 TCM, across patients, and observed that cell types were indeed strongly correlated in their response magnitude (Fig. 3d). Therefore, CPD provided the added insight that reduced IL-10 response is associated with an increase in the baseline level of pSTAT3 and is coordinated across cell types, varying in magnitude across patients according to their disease cohort.

**Fig. 3.**
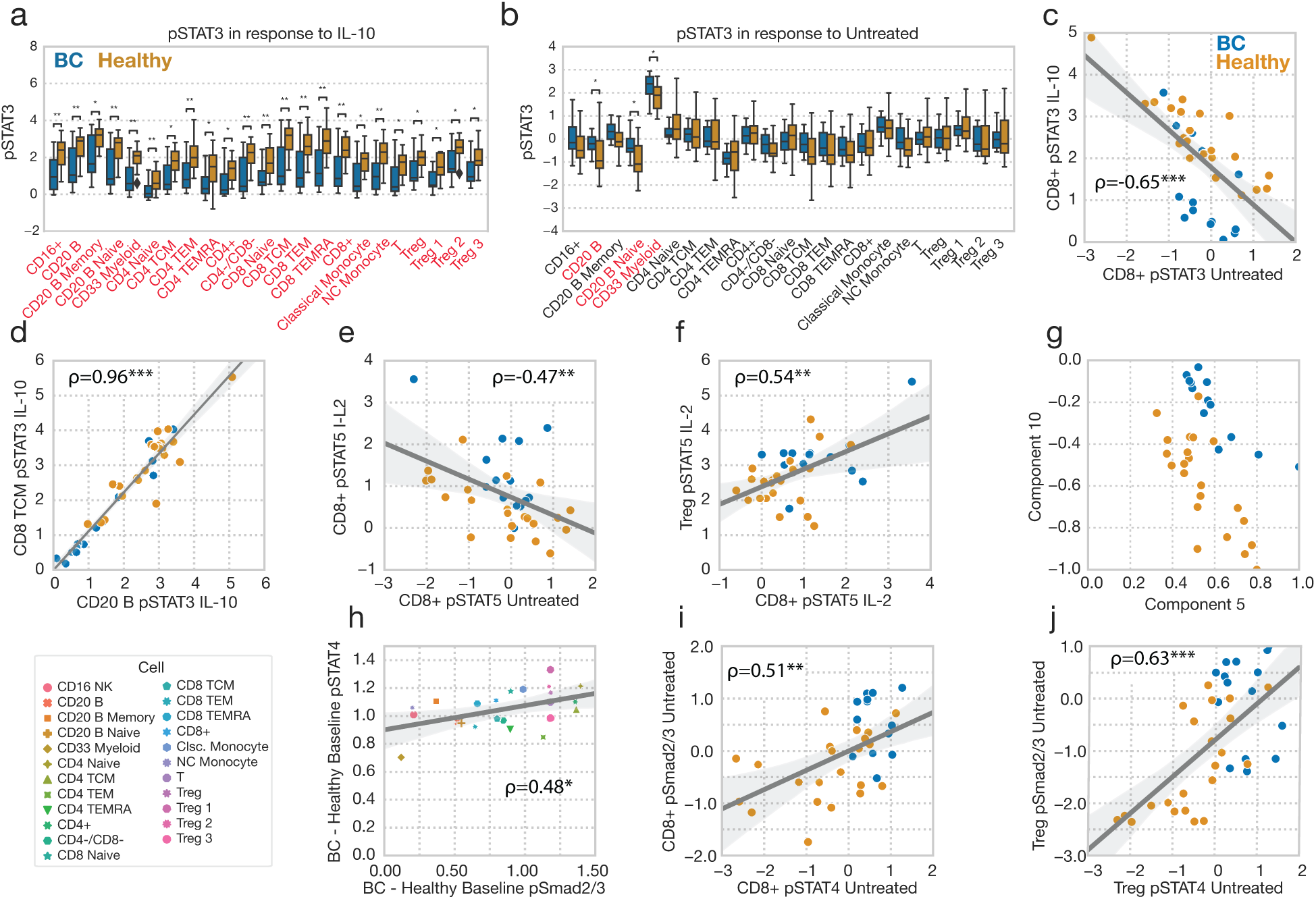
Immune dysregulation is more prominently reflected in patterns of signaling response changes. **(a)** STAT3 phosphorylation response to IL-10 across cell types, grouped by patient status. Cell types with statistically significant differences between BC and healthy status are colored red. **(b)** Baseline pSTAT3 in untreated cells across cell types, grouped by patient status. **(c)** Baseline pSTAT3 versus IL-10-induced pSTAT3 in CD8-positive cells, across patients. **(d)** pSTAT3 response to IL-10 across patients in B cells versus CD8 TCM. **(e)** Baseline untreated versus IL-2-induced STAT5 phosphorylation in CD8-positive cells. **(f)** IL-2-induced pSTAT5 in CD8-positive cells and Tregs. **(g)** Components 4 and 5 from the CPD factorization in Figure 2. **(h)** The difference in average Smad1/2 phosphorylation between the BC and healthy cohorts versus the same quantity for STAT4 phosphorylation, plotted for each cell type. **(i–j)** Baseline pSmad2/3 versus pSTAT4 across patients in CD8-positive cells (i) and Tregs (j).

Second, we observed that the patterns of signaling dysregulation associated with BC more effectively than individual measurements alone. For instance, IL-2 response was increased within CD8 cells in BC patients. However, using baseline versus IL-2-induced pSTAT5 (Fig. 3e), or CD8 versus T_reg_ responses (Fig. 3f), incompletely separated BC from healthy patients. By contrast, the pSTAT5-associated component 5 led to almost perfect separation when combined with component 10, summarizing the baseline pSmad2/3 levels (Fig. 3g). Through this, we observed that CPD, though unsupervised, can better define patient groups by defining integrative signatures of response.

We found that CPD-defined factors could additionally identify coordinated changes across pathways. Component 10 represents an increase in basal Smad1/2 and STAT4 phosphorylation in untreated cells across most cell types. While a per-measurement analysis reveals these same differences, the CPD results reveal that these changes are coordinated in the magnitude of the difference across cell types (Fig. 3h) and patients (Fig. 3i/j). Therefore, profiling and analyzing these differences in a multidimensional way reveals how the changes are shared across dimensions and improves our ability to resolve these differences.

### CPD identifies coordinated patterns of PBMC receptor abundance variation

Cell surface receptors delineate cell types and which cells are capable of cytokine and growth factor responses^23^. Therefore, we next investigated whether receptor abundance itself may associate with cancer status or define the differences in cytokine response.

For instance, reduced IL-6 signaling in BC patient PBMCs, associated with poor clinical prognosis, has correlated with reduced IL-6 receptor expression^6^. In a separate sample, we measured receptor abundance profiles in each patient and cell type across the patient cohort. Cells were stained for the cognate receptors of each cytokine stimulant (e.g., IL10R, TGFβRII), receptors with signaling pathways converging on our cohort of measured transcription factors (e.g., IL12RII–pSTAT4, IL7Ra–pSTAT3/5), as well as the checkpoint proteins PD-1 and PD-L1.

These data can be naturally organized into a three-dimensional tensor, with patient, cell type, and receptor dimensions. We again summarized and visualized these data using tensor decomposition (Fig. 4a). Five components explained over 60% of the variance in our dataset (Fig. 4b). As with the cytokine response data, tensor factorization reduced the dataset more efficiently than PCA (Fig. 4c). We again validated our CPD approach for the receptor dataset by testing the model’s ability to impute missing values (Fig. S2). The five-component model was selected based on its optimal association with BC disease (Fig. 4d). Logistic regression identified that component 2 almost perfectly distinguishes the breast cancer and healthy patients, with higher values associated with cancer (Fig. 4e). Still, hierarchical clustering of the patient factors broke both the healthy and cancer patients into two groups each, reflecting that patient-to-patient variation beyond just that contained in the highly BC-specific component 2 was present (Fig. 4f).

**Fig. 4.**
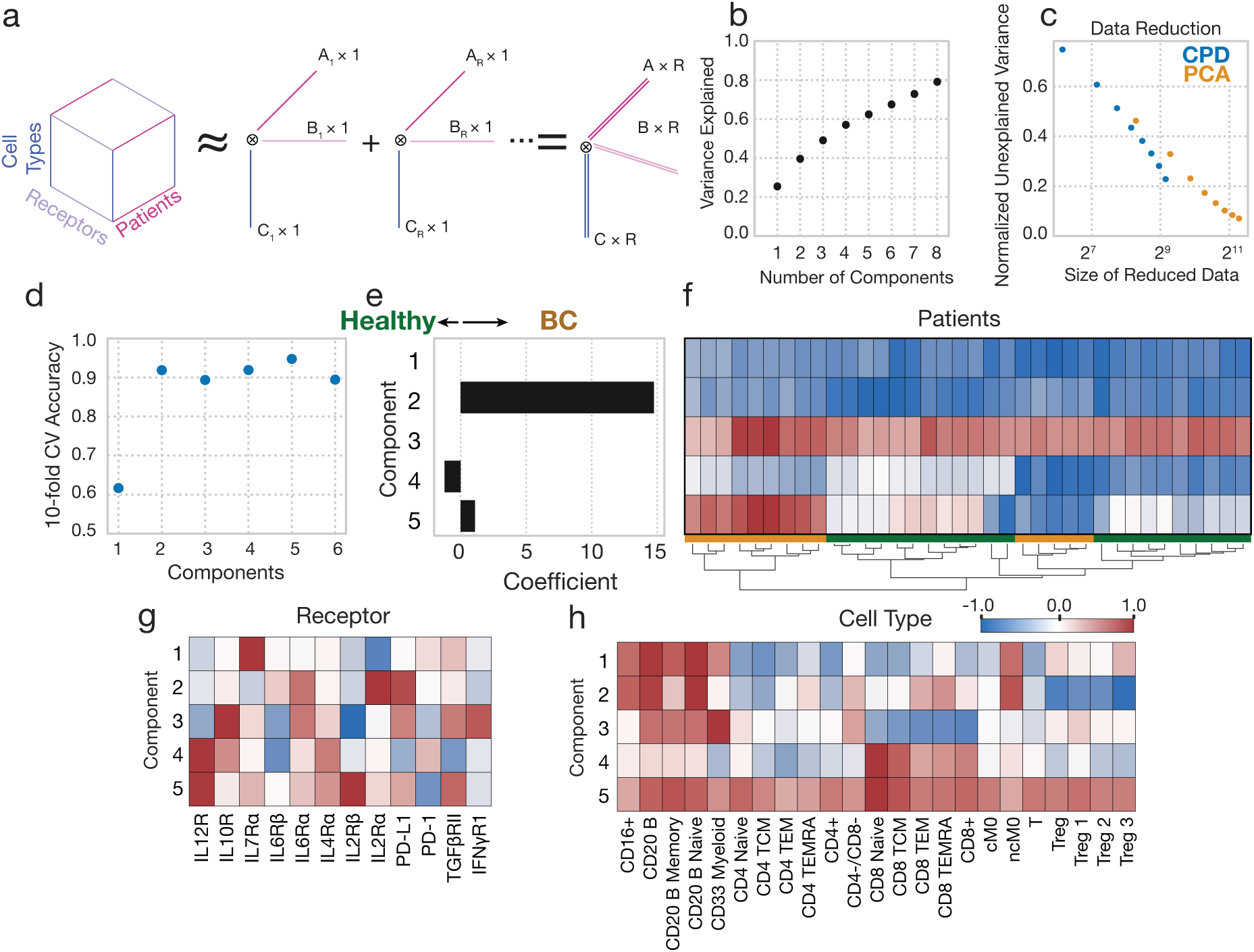
CPD reveals patterns of receptor abundance variation. **(a)** Schematic of CPD. Receptor data is organized into a three-dimensional tensor with axes of patient, cell type, and receptor. This tensor is reduced into sum of the outer products of rank 1 tensors (components), allowing for easy visualization. **(b)** Percent variance reconstructed (R2X) versus the number of components used. **(c)** The remaining error on reconstruction, normalized to the total dataset variance, versus the size of the dataset after decomposition using CPD or PCA. **(d)** The accuracy of a logistic regression classifier upon 10-fold cross-validation, using the CPD patient factors with varying numbers of components. **(e)** The weights associated with each component for a logistic regression classifier fit to patient factors using a 5-component decomposition (healthy = 0, BC = 1). **(f)** A heatmap of the patient factor matrix, with the patients hierarchically clustered. Patient status is indicated by the coloring along the bottom. **(g– h)** Component values for each receptor (g) and cell type (h).

As before, one can trace a single component across the factor matrices to make inferences about that pattern within the dataset. Component 2 represents variation in IL2Rα, IL6Rα, and PD-L1 across all samples and increased in BC (Fig. 4g), with increased abundance in CD8 T cells, non-classical monocytes, and B cells, alongside decreased abundance in (T_reg_s; Fig. 4h). This alteration was consistent among T_reg_ subsets I, II, and III, each of which have been shown to play distinct roles in immunosuppression and correlate in distinct manners with breast cancer outcome^8,24^. We observed that variation in several receptors was summarized by single components, such as the three receptors associated with component 2, indicating correlations in these receptors’ expression across patients and cell types (Fig. 4g). Thus, overall, CPD succinctly summarized these patterns of receptor variation across both cell types and patients.

### Dissecting the patterns of receptor variation suggest concerted molecular programs

Using the CPD results as a guide, we further dissected the patterns of receptor variation. Interestingly, we did not see levels of PD-1 in CD8 T cells as a distinguishing feature of BC, nor did we see significant changes in PD-1 abundance in other CD8 subsets (Fig. 5a). Component 2 identified most of the cell types and receptors with highly significant differences on a per-measurement basis (Fig. S6). PD-L1 was elevated in TCM, TEM, and TEMRA-positive CD8 cells, though not in the naïve subset, and was upregulated in several B cell subsets (Fig. 5b). IL6Rα was also found in higher amounts in roughly the same cell subsets (Fig. 5c). IL2Rɑ, by contrast, decreased with BC in regulatory T cell subsets (Fig. 5d), consistent with the cell populations’ negative weighting on component 2 (Fig. 4h).

**Fig. 5.**
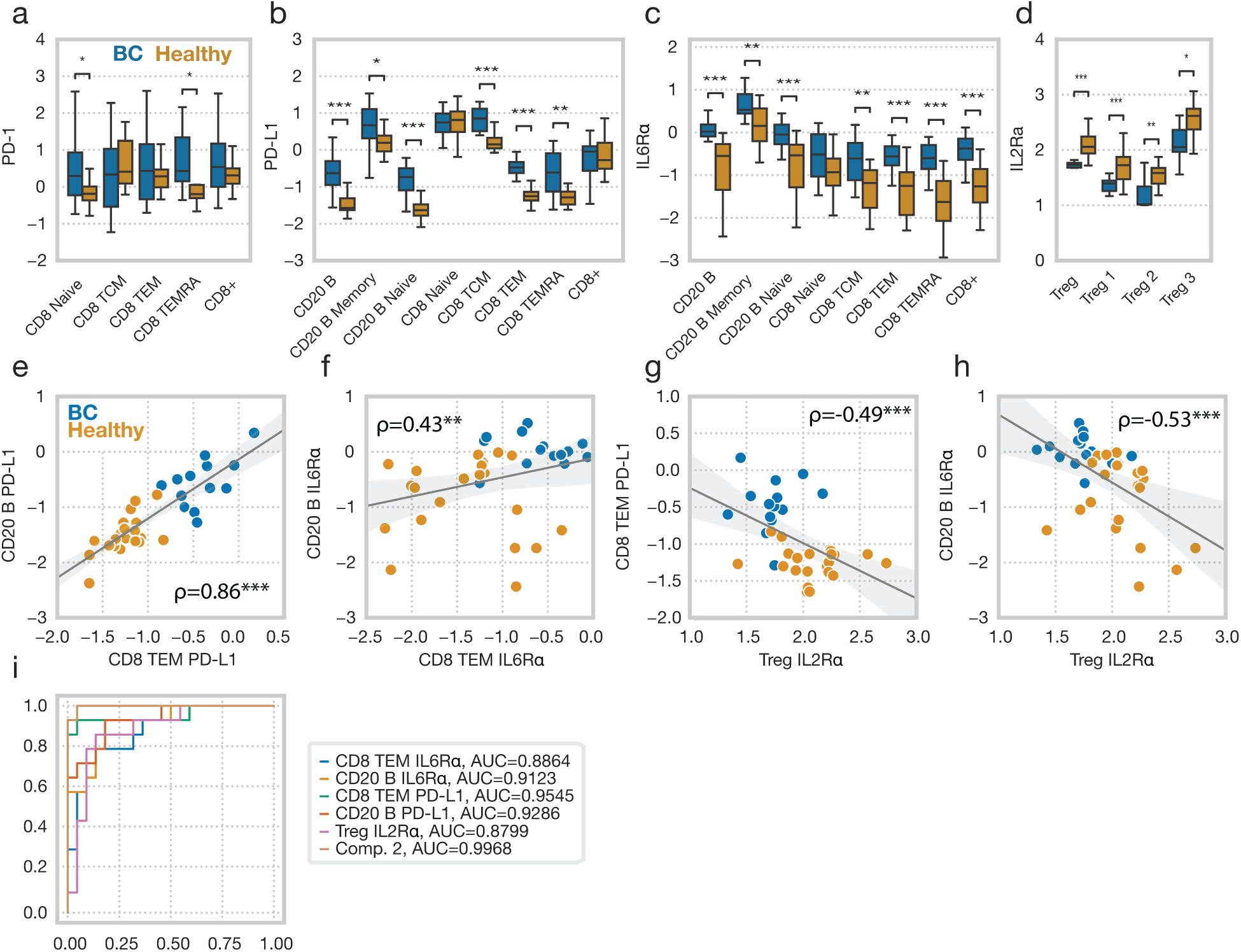
Dissecting the patterns of receptor variation suggest concerted molecular programs. **(a)** Mean relative abundance of PD-1 across CD8-positive cell types, grouped by patient disease status. **(b)** Mean relative abundance of PD-L1 across CD8-positive and B cells, grouped by patient disease status. **(c)** Mean relative abundance of IL6Rα across CD8-positive and B cells, grouped by patient disease status. **(d)** Mean relative abundance of IL2Rɑ in Tregs, grouped by patient disease status. **(e–h)** Correlation across patients between CD8^+^ TEM PD-L1 and CD20 B PD-L1 (e), CD8^+^ IL6Rα and CD20 B IL6Rα (f), Treg IL2Rɑ and CD8 TEM PD-L1 (g), and Treg IL2Rɑ and CD20 B IL6Rα (h). (i) ROC curve for the separation provided by several of the receptor amounts measurements, alongside component 2. *, **, and *** represent p-values less than 0.05, 0.005, and 0.0005, respectively.

Given that IL2Rɑ differences were represented by the same component in the CPD analysis but showed up in univariate analysis within a separate cell subpopulation than the IL6Rα and PD-L1 differences, we wondered whether a per-measurement view masked additional relationships among these measurements. Each component in CPD represents variation shared among patients, receptors, and cell types. Therefore, we explored whether we could observe these coordinated changes. Indeed, we observed that PD-L1 abundance was strongly correlated among patients between CD8 TEM and B cells (Fig. 5e), as was IL6Rα amount (Fig. 5f). Furthermore, the pattern of coordinated changes extended to comparisons across both receptors and cell types. Plotting the abundance of IL2Rɑ among Tregs against PD-L1 in CD8 TEM cells, we observed a correlation across patients (Fig. 5g). As another example, we observed a correlation between IL2Rɑ abundance in Tregs and IL6Rɑ abundance in B cells (Fig. 5h). These results show that the CPD-defined components indeed represent coordinated changes within and across cytokine pathways.

Lastly, we surmised that the coordinated changes across receptors and cell types might better distinguish BC from healthy patients either due to averaging over measurement noise or capturing inherently multivariate patterns. Indeed, when comparing the classification ROC of component 2 versus B cell or CD8 TEM IL6Rα or PD-L1, as well as Treg IL2Rɑ, the CPD component still led to the optimal separation of patient status (Fig. 5i). Thus, in total, the receptor changes we observed represent a coordinated pattern of immunologic reprogramming across cell types and receptors.

### Receptor abundance defines cell type- but not BC-specific responses

We then sought to examine whether receptor-level differences in cytokine signaling regulation can explain the differences in basal and induced signaling in BC patients. As expected, receptor expression clearly defined which cell types respond to certain ligands. For instance, average IFNγ-induced STAT1 phosphorylation across cell types correlated with IFNγR1 abundance (Fig. 6a). IL2Rɑ abundance also correlated with which T cell subsets were most responsive to IL-2 (Fig. 6b). Thus, we wondered if patient-to-patient differences in response could also be explained by variation in a cytokine’s cognate receptors.

**Fig. 6.**
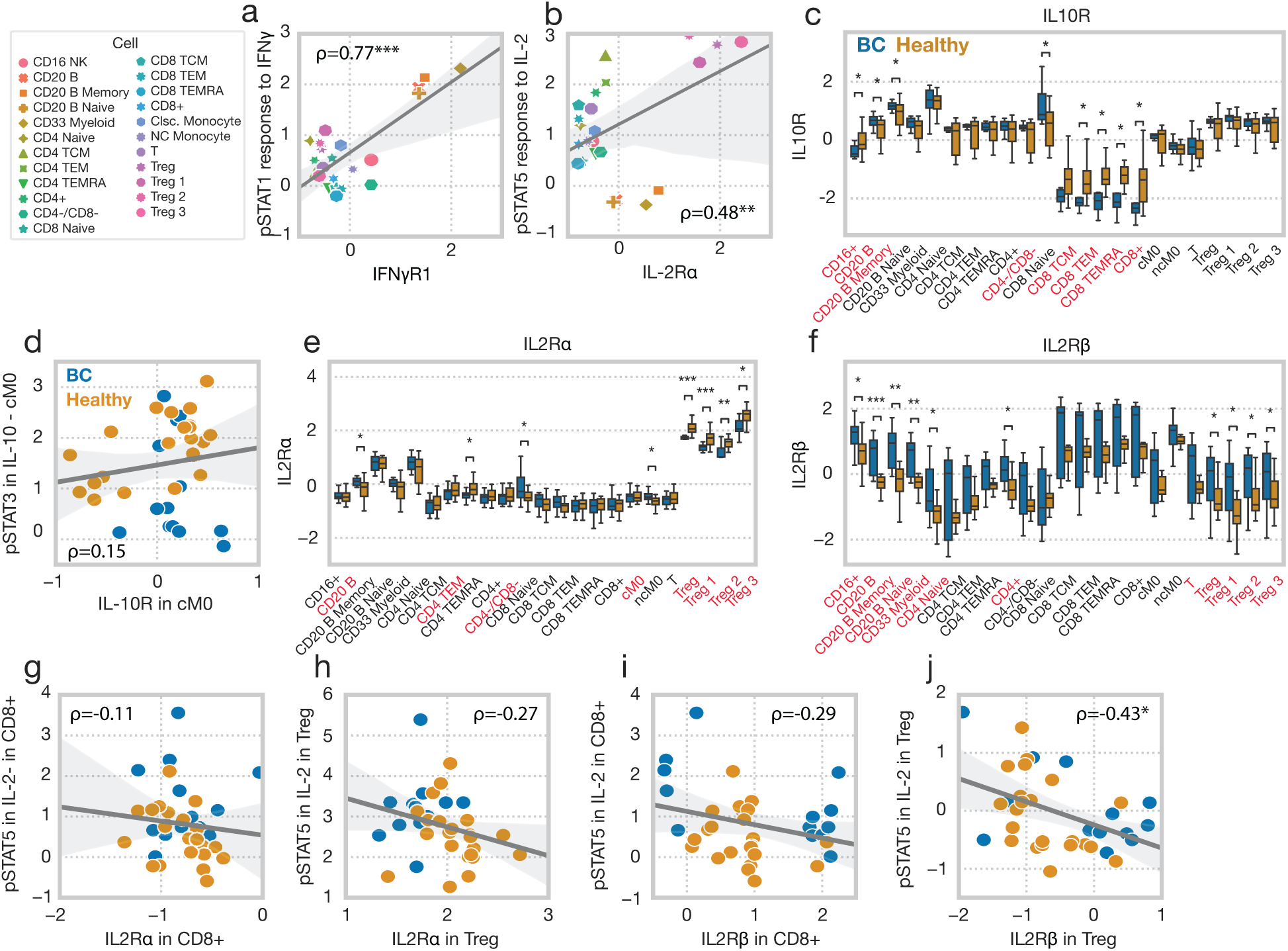
Receptor abundance defines cell type- but not BC-specific responses. **(a)** Patient-average IFN-induced STAT1 phosphorylation versus IFNγR1 abundance across cell types. Induced responses are reported as z-scored delta MFI. **(b)** Patient-average IL-2-induced pSTAT5 versus IL2Rɑ abundance across cell types. **(c)** IL10R abundance across cell types, grouped by patient status. **(d)** IL-10-induced pSTAT3 versus IL10R abundance in classical monocytes, across patients. Cell types which feature statistically significant responses between BC and healthy status are listed in red. **(e)** IL2Rɑ abundance across cell types, grouped by patient status. **(f)** IL2Rβ abundance across cell types, grouped by patient status. **(g–j)** IL-2-induced pSTAT5 versus IL2Rɑ (g/h) or IL2Rβ (i/j) in CD8^+^ cells (g/i) or Tregs (h/j).

One of the most prominent changes we observed was a reduced response to IL-10 in BC, and so we first examined whether IL10R was expressed at a lower level in BC. Although IL-10 response was reduced across all cell types, we only observed a reduction in IL10R abundance within CD8 cells and B cells (Fig. 6c). Consequently, while IL-10 responses were significantly reduced in classical monocytes, there was no correlation between blunted responses and the levels of IL10R (Fig. 6d).

Similarly, while IL-2 responses were uniformly increased across cell populations in BC, IL2Rɑ was unchanged in most cell types, and actually decreased in Tregs (Fig. 6e). IL2Rβ was heterogeneously expressed across BC patients, such that it was increased in a subset of patients (Fig. 6f). However, because of this heterogeneity, neither IL2Rɑ or IL2Rβ correlated with the differences in IL-2 response in CD8+ cells or Tregs (Fig. 6g–j).

Differences in receptor amounts also did not have a 1-to-1 correspondence with baseline pathway activation. The general increase in basal pSTAT4 was not explained by IL12RI levels, which were generally unchanged (Fig. S6l). TGFβ-RII was upregulated in CD8^+^ cells, which may partially explain their increased basal pSmad2/3 levels, but not the increases in other cells (Fig. S6b). While the IL6Rα was increased in BC across many populations (Fig. S6h), we did not observe an increase in BC-associated IL-6-induced pSTAT3 response, but rather decreased responses (Fig. S5c). Thus, in total, we found that the presence of a cytokine’s cognate receptor defined the population-specificity of a response, but not variation in response across patients.

### Patterns of coordinated receptor-signaling dysregulation suggest mechanisms of response reprogramming both globally and in a cell type-specific manner

Given that receptor abundance did not consistently inform the basis of the BC-associated immune signaling differences, we next wondered whether the receptor levels and basal / induced signaling associated with each other in unintuitive ways.

First, to explore these associations in a cell type- and measurement-specific manner, we calculated the partial correlation across each significantly different induced or basal phosphorylation level as well as each differentially abundant receptor and used hierarchical clustering to examine the correlation patterns (Fig. 7a). In CD8^+^ cells, we observed groups of strongly correlated features: pSTAT3 response to IL-10 was strongly correlated with pSTAT5 response to IL-2, and both changes were significantly anticorrelated with increases in IL6Rα, basal pSmad2/3, IL-12RI, and IL2Rβ. Another highly correlated module involves basal pSTAT4, PD-L1, TGFBII, and IL10R. While these associations were similar in CD4^+^ cells, inconsistencies of the exact patterns present in each cell type, as well as the magnitude of those correlations made it difficult to compare patterns across cell types (Fig. 7b). Thus, a more global view of how patterns related across cell types and patients was required.

**Fig. 7.**
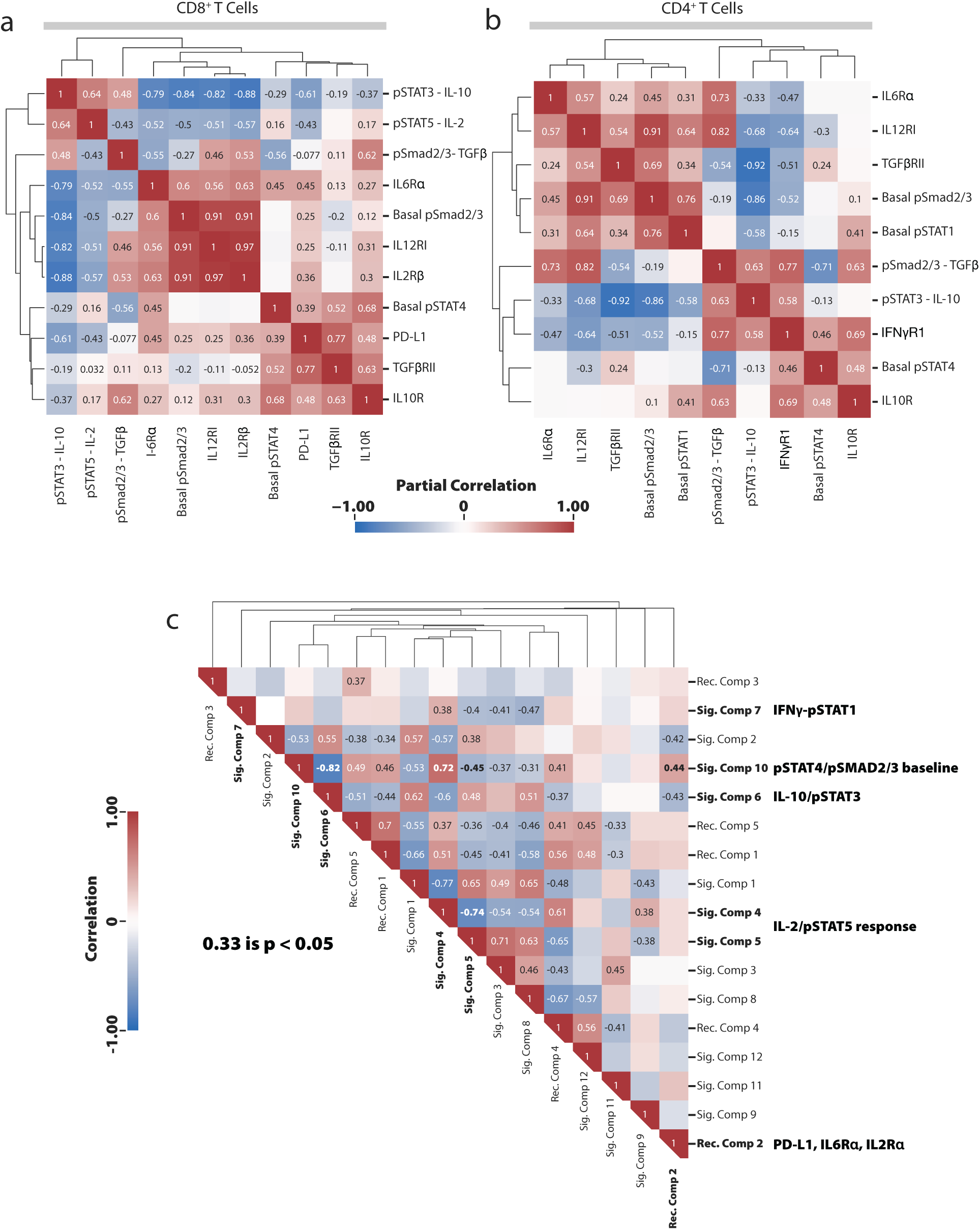
Patterns of coordinated receptor-signaling dysregulation suggest mechanisms of response reprogramming both globally and in a cell type-specific manner. **(a–b)** Partial correlations among measurements with statistically significant differences in CD8^+^ (a) or CD4^+^ (b) T cells. Correlations are calculated across patients, specifically within the BC cohort. **(c)** Correlations across all patients among the receptor and signaling component patterns.

Given that the CPD-derived patterns summarized the variation we observed across all other dimensions, we then used them to identify correlations in patterns of patient-to-patient variation on a more global scale. The directionality of each component pattern can vary depending on the other factors and so we clustered correlations according to their absolute value, building a similar clustering (Fig. 6c). The dominant BC-associated pattern of receptor expression we observed (increased and IL6Rα and PD-L1 in CD8^+^ and B cells, increased IL2Rα; component 2, Fig. 4) were significantly correlated with the pan-cell type increases in pSmad2/3 baseline and decreased IL-10-induced pSTAT3 (component 6 & 10, Fig. 2). Additionally, IL-2-induced as well as basal pSTAT5 (component 4 & 5, Fig. 2) increases were also strongly correlated with these alterations in signaling and receptor expression. These data suggest that, while each of these changes manifested within a different series of the profiling measurements, they were coordinated in their presentation across patients, and so could be expected to underly a common pathway of dysregulation. Notably, many other complex relationships between non-BC associated patterns were additionally identified, painting a picture of highly coregulated immune alterations.

## DISCUSSION

Here, we systematically dissected the altered cytokine signaling responses found in breast cancer patients as compared to healthy controls. Our approach directly accounts for the multidimensional nature of cytokine response, which varies according to stimulation, cell type, and the measured signaling product, as well as on a per-patient basis. Accordingly, we profiled cytokine response across these dimensions and found that the responses of PBMCs in breast cancer are altered within and across cell types, cues, and pathways. We also observed patterns of changes in receptor abundance which were coordinated between cell types. However, while we observed that variation in receptor abundance was associated with response to the cognate cytokine across cell types, it did not directly explain differential response between patients. Patterns of dysregulated receptor abundance instead correlated with alterations in signaling pathways outside of their cognate pathways. Thus, we found that cytokine response dysregulation occurs through coordinated programs.

A central finding of our work is that there is a coordinated disruption in the cytokine responses of PBMCs in breast cancer patients, compared to healthy donors, across several pathways, cytokines, and cell types. Both high basal levels and reduced responsiveness of Smad1/2 to TGFβ were associated with ER+ BC and with suppression of STAT3 phosphorylation, at baseline and in response to IL-10. Additionally, this pattern was correlated among patients with reductions in IL10R, IL12R, and IL4Rα in B cells and CD8^+^ T cells. This result was particularly interesting as IL-10 is known to drive immunosuppression in most populations, but also more recently been shown to be a key driver of CD8^+^ T cells’ anti-tumor activity^25,26^. Heightened pSmad2/3 through increased TGFβ has been shown to drive cytotoxic cell exhaustion^27^. Likewise, pSTAT4, which has been shown to be associated with increased IL-6 production, was found at significantly elevated levels in unperturbed BC immune cells^28^. We also observed that increased PD-L1 expression was strongly correlated with TGFβ-RII expression suggesting that these two immunosuppressive proteins work in tandem in CD8^+^ cells to suppress effector response. Interestingly, we found that IL6Rα expression was negatively correlated with pSTAT3 response to IL-10, suggesting that heightened IL6Rα and IL-6 in cancer patients may be dominating STAT3 signaling and in turn suppressing the IL-10 responsiveness. In B cells, in addition to increased basal pSTAT4 and pSmad2/3, we found reductions in IL-10 responsiveness through pSTAT3, increased IFNγ responsiveness through pSTAT1, and increased PD-L1, IL6Rα, and IL2Rβ expression were associated with BC status. Lower pSTAT3 responsiveness to IL-10 potentially reflects potentiation after exposure to the cytokine, and positive staining for PD-L1 is a hallmark of IL-10-secreting regulatory B cells which promote tolerance in allergy and autoimmunity and have previously been associated with invasive breast cancer^29^. Finally, several patterns associated with Tregs were also found to be informative of BC disease status, such as decreased IL2Rα, and increased responsiveness to IL-2 through pSTAT5. These results are intriguing and at first glance contradictory, as IL2Rα potentiates Tregs sensitivity to IL-2^11^. These results may point to Tregs with low activation due to IL-2 starvation, as other B and T cell subsets were found to express high amounts of IL2Rβ (Fig. S6)^30^. A full table of features found to define BC patient features was compiled in Table 1.

**Table 1.** Full table of patterns associated with Breast Cancer.

| **Pattern** | **Cell Types** |
| --- | --- |
| Decreased STAT3 phosphorylation in response to IL-10 | Pan-cell type, and particularly in CD8<sup>+</sup> cells |
| Increase STAT5 phosphorylation in response to IL-2 | Primarily in T<sub>reg</sub>s |
| Increased basal pSTAT4 | Pan-cell type, and particularly in CD8<sup>+</sup> cells |
| Increased basal pSmad2/3 | Pan-cell type, and particularly in CD8<sup>+</sup> cells |
| Increased STAT1 phosphorylation in response to IFNγ | B and CD33<sup>+</sup> myeloid cells |
| Increased PD-L1 abundance | B and CD8<sup>+</sup> effector cells |
| Increased IL2Rα abundance | T<sub>reg</sub> cells, CD4<sup>+</sup> cells |
| Increased IL6Rα abundance | CD16<sup>+</sup>, B, CD8<sup>+</sup> cells, and monocytes |
| Increased IL2RB abundance | B cells and CD8<sup>+</sup> cells |
| Decreased IFNγR1 abundance | CD16<sup>+</sup> cells, CD4<sup>+</sup> cells |

Taken together, our results point to an overall shift in immunologic features commonly associated with dysregulated immune responses in other contexts. Elevated pSMAD2/3 and induction with TGFβ, combined with elevated pSTAT4, is suggestive of a Th17-like response^31^. Indeed, Th17 cells are found in greater numbers within breast tumors, and another inducer of Th17 responses, IL-6, is elevated in a variety of cancers, playing a central role in cachexia^32–34^. Finally, we found that IL2Rα was found in decreased abundance in the Tregs of breast cancer patients, a finding which has been reported in a broad panel of autoimmune diseases^35^. Our analysis also revealed several patient features that correlate with more traditionally cancer-associated immunosuppressive features, which were like those found in the suppression of autoimmune diseases. For example. We found that B cells displaying B_reg_-like phenotypes were selectively found in BC patients. These cells have previously been found to be important to the resolution of autoimmune diseases such as rheumatoid arthritis and experimental autoimmune encephalomyelitis^36,37^. Furthermore, we also found increased PD-L1^hi^ CD8^+^ cells in BC patients with reduced capacity to respond to IL-10. These features are typical of regulatory CD8^+^s, a cell type previously reported as resident in breast cancer tumors and with important roles in the control of systemic lupus erythamatosis^38,39^. Thus, and is reminiscent of systems where inflammatory features are only held in check by compensatory increases in several immunosuppressive cell types and signaling processes, a state which resembles that of with suppressed autoimmune responses.

An important observation with of our work is that, while most characterization of tumor immunity has focused on the local microenvironment or lymph nodes, our observations were made in peripheral cells with no selection for antigen-specific cells^40^. Furthermore, we find that several changes in cytokine responsiveness and marker expression previously associated with breast cancer are in fact coordinated changes, which could be the result of a shared process of tumor-driven reprogramming^41^. Previous work has shown that these global changes are not explained by changes in cytokine abundance in the blood^7^, and thus may be a result of immunologic reprogramming within lymph nodes^42^, or reprogramming during local trafficking of cells through the tumor.

While we were able to establish differences in cytokine response and baseline receptor abundance, these data establish a strong basis for a more in-depth characterization of the global immunologic differences that develop with cancer. Cancers develop through a progressive process of immunoediting and then escape from immune control^43^.

Analyzing the changes in cytokine responsiveness and receptor abundance we identified here, alongside other forms of profiling information such as transcriptional and epigenomic, will help to reveal both how these cytokine pathway changes arise and how they are linked to functional changes within immune populations^44,45^. While breast cancer has been poorly responsive to immunotherapy, given the significant receptor expression and cytokine response changes we observe, there should be clear and informative transcriptional changes within peripheral immune cells.

An open question is also how these changes within the periphery reflect changes within other immune sites, such as lymph nodes and the tumor microenvironment^8,40^. One might expect that through circulation peripheral cells reflect changes in the local tumor microenvironment^8^, or that differential effects in trafficking mean that the periphery reflects the absence of certain cytokine-responsive cells, or presence of cells which stymie response^46^. Finally, while our previous work has shown that dysregulated signaling response correlates with worsened rates of relapse^6^, profiling these coordinated changes in other patient populations that vary in outcomes and treatment response, along with identifying other features correlated with cytokine responsiveness, may allow us to more fully assess how these changes might provide a useful prognostic readout of immune functionality.

Our work demonstrates the value of two methodologic approaches. First, profiling and analyzing cell perturbations such as cytokine responses in a multi-dimensional manner can reveal coordinated changes across cell types. Specifically, we identified changes in response that are localized to baseline levels of signaling, changes localized to cell types, and global changes across cell types. Restricting any one of the dimensions we explored—patients, cell types, cytokines, or responsive proteins and markers—would have consequently restricted our conclusions. Second, including healthy controls provided a baseline for immune response from which we could contrast the responses observed in cancer samples. Breast cancer, like so many malignancies, demonstrates heterogeneity both within and between patients. However, this work shows that there are consistent and substantial changes across patients, potentially reflecting concerted tumor-directed reprogramming that is critical for disease progression and response to therapy.

## MATERIALS AND METHODS EXPERIMENTAL METHODS

### Human samples

Peripheral blood samples were obtained from consented patients (IRB #21368 and #19186) with ER+ breast cancer at City of Hope. Patient characteristics are summarized in Supplemental Table S1. All patients who consented to this study had no previous history of breast cancer and were estrogen receptor-positive (ER+), HER2/neu negative (HER2-). Patient’s blood was drawn into EDTA-containing tubes. Peripheral blood mononuclear cells were isolated by Ficoll-Paque (Cytiva, Marlborough, MA, USA) density centrifugation following the manufacturer’s protocol and cryopreserved in 10% DMSO FBS. Age-matched healthy control peripheral blood samples were obtained from City of Hope Blood Donor Center.

### Cell culture

Cryopreserved PBMCs were thawed and rested overnight (16h) in RPMI 1640 medium supplemented with 10% fetal bovine serum, 1% penicillin-streptomycin-glutamate (PSG) at 37℃, 5% CO_2_. PBMCs were counted using a hemocytometer, and viability was determined by trypan blue exclusion (Sigma-Aldrich). Subsequently, cells were stimulated in 96 deep-well V plates at a concentration of 0.5–1 × 10^6^ cells/ml in fresh RPMI 1640 medium (Thermo Fisher Scientific Inc., MA) supplemented.

### Cell signaling

After resting period, PBMCs were either untreated or stimulated with IFNγ (50 ng/ml), IL-10 (50 ng/ml), IL-6 (50 ng/ml), IL-4 (50 ng/ml), or TGFβ (50 ng/ml) (Peprotech, Rocky Hills, NJ, USA) at 37℃ for 15 min, followed by fixation with 1.5% paraformaldehyde (PFA) for 10 min at room temperature. Following PFA fixation, cells were washed with PBS 1x and permeabilized using ice-cold 100% methanol. Following methanol fixation, cells were stored at −80℃. Cells were then washed three times with staining buffer (PBS supplemented with 1% FBS) before antibody staining.

### Phospho Flow Cytometry

The following antibodies were used: STAT4-AF647 (clone 38/p-Stat4), CD14-APC-Cy7 (clone HCD14), CD20-AF700 (clone H1), STAT6-V450 (clone 18/pStat6), PD-L1-BV510 (clone 29E.2A3), CD3-BV570 (clone UCHT1), PD1-BV605 (clone EH12.1), CD33-BV750 (clone p67.6), CD27-BV786 (clone L128), CD45RA-BUV395 (clone HI100), CD4-BUV563 (clone SK3), CD16-BUV737 (clone 3G8), CD8-BUV805 (clone SK1), STAT3-

AF488 (clone 4/p-Stat3), STAT1-Percp-Cy5.5 (clone 4a), SMAD2/3-PE (clone O72-670), Foxp3-PE-CF594 (clone 259D/C7), STAT5-PE-Cy7(clone 47). Dilutions of antibodies were prepared based on the manufacturer’s recommendations and optimized for staining conditions in preliminary experiments. Incubation was carried out for 45 minutes at room temperature. All antibodies used were purchased from Biolegend San Diego, CA, USA) or BD Biosciences (Franklin Lakes, NJ, USA)

### Cytokine receptor flow cytometry

The following antibodies were used: CD45RA-BUV395 (clone HI100), CD3-APC-Cy7 (clone UCHT1), CD16-BUV737 (3G8), CD16-APC-Fire810 (3G8) CD119 (IFNγR1)-BB660 (clone GIR-208), CD33-BV750 (clone p67.6), CD27-AF700 (clone O323), CD14-BUV496 (clone MOP9), CD132 (IL-2Rγ)-BB700 (clone TUGh4), CD210 (IL-10R)-PE-Cy7 (clone 3F9), CD122 (IL-2Rβ)-BV786 (clone MIKB3), TGF-βR2-APC (clone W17055E), PD-L1-BV510 (clone 29E.2A3), CD212 (IL-12R1)-BV421 (clone 2.4e6), PD1-BV605 (clone EH12.1), CD8-BUV805 (clone SK1), CD4-BUV563 (clone SK3), CD25-BB515 (clone MA251), CD124 (IL-4Rα)-BB630 (clone HIL4R-M57), CD124 (IL-4Rα)-PE (clone HIL4R-M57), CD130 (IL-6Rβ)-PE-CF594 (clone M5), CD130 (IL-6Rβ)-BUV737 (clone M5), CD127-BV650 (clone A019B7). Dilutions of antibodies were prepared based on the manufacturer’s recommendations and optimized for staining conditions in preliminary experiments. Incubation was carried out for 30 minutes at 4°C. All antibodies used were purchased from Biolegend San Diego, CA, USA) or BD Biosciences (Franklin Lakes, NJ, USA).

### Data acquisition and gating strategy

Stained cells were analyzed using the Cytek Aurora flow cytometer with a 355 nm, 405 nm, 488 nm, 561 nm, and 640 nm laser configuration. Compensations were established using single-stained controls and a negative control sample. A total of 50,000 to 100,000 events were acquired at a data rate of 1000 events/s. Cell populations were gated as shown in supplementary figures S1.

## MODELING

### Gating and Preprocessing

For all quantification of cellular species abundances, whether cell type markers, signaling products, or receptor amounts, the mean fluorescent intensity (MFI) of flow cytometry data was calculated to determine signal. Cell population gating was performed as shown in figure S1. Prior to decomposition, the mean signaling response data was background subtracted on a per-patient, per-cell type basis, and normalized for each signaling marker according to the maximum signal observed for that marker. Receptor data was normalized by z-scoring each receptor’s signal.

### Canonical Polyadic Decomposition

Before analysis, we reorganized our measurements into a third- or fourth-order tensor, or a three- or four-dimensional array with axes representing parameters over which the profiling was conducted. Within the CP decomposition model, a tensor is described as a sum of the tensor product (⊗) of rank-one components that represent the contribution of each mode. For instance, in the three-mode case:

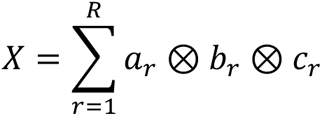

Where *a_r_*, *b_r_*, and *c_r_* are the r^th^ column of the factor matrices *A*, *B*, and *C*, which overall summarize how each pattern is represented across the three dimensions.

While many algorithms exist for deriving CP factorizations, we applied alternating least squares (ALS), wherein each mode is iteratively solved using least squares. As an iterative procedure, ALS must be initialized with a starting estimate of the factorization.

The CPD decomposition was initialized using the right-hand *r* eigenvectors from SVD decomposition of the tensor flattened along each mode.

ALS applies the observation that, given factor matrices for two of the modes (e.g., modes 2 and 3), the optimal factor matrix for the remaining mode can be solved for as the least squares solution between the tensor unfolding and Khatri-Rao product of the known factors:

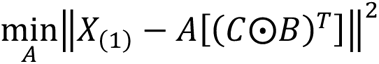

*X*_($)_ represents the tensor unfolding of *X* along mode 1, and *C*⊙*B* represents the Khatri-Rao product of *C* and *B*. After solving for *A*, ALS proceeds for each of the remaining modes, completing one iteration of the algorithm by building a representation of the other factors using the Khatri-Rao product, and then applying least squares to solve for the left-out factor matrix.

In addition to the ALS scheme described above, we applied the line search routine described by Bro^47^. Briefly, after two rounds of ALS, the difference between the last two fitting iterations was used to look ahead by a line search step equal to √^!^ *N*, where *N* is the iteration number and *l* is the line search quotient, initially equal to 2. If the error of the line search-updated factorization is less than the normal ALS update, then the line search result was accepted. Otherwise, the ALS result was used. After four straight rejections the line search quotient was increased by 1 to reduce the line search distance.

### Censored Alternating Least Squares

The censored least squares algorithm is solved similarly to ALS but differs in its approach of handling the missing value problem. Rather than imputing missing values, we first grouped variables during the least-squares solve based on their pattern of missing values within the dataset. Solving was then performed on each group, with the missing rows removed during each part of the solving process. In this way, the least squares solution was solved without inclusion of the missing values or the need for imputation during fitting.

### Logistic Regression

Regularized logistic regression was implemented using the implementation provided by scikit-learn, with an l1 penalty^48^. Solving was performed with the Stochastic Average Gradient (SAGA) solver^49^, a maximum iteration number of 5,000 and tolerance of 1.0 × 10^*+^. The regularization strength was determined through cross-validation using the LogisticRegressionCV and the default parameters regarding attempted regularization strengths. We applied repeated, stratified, 10-fold cross-validation, with 20 repeats throughout the analysis.

### Quantification and statistical analysis

For each figure, descriptions of pertinent statistical analyses or metrics used, the number of replicates of experiments performed, and the values of confidence intervals can be found in its corresponding figure caption. n indicates the number of times a particular experiment was performed (duplicate, triplicate, etc.) within each figure.

To test for statistical differences between induced and basal signaling activation, as well as receptor abundance, a Mann-Whitney U was performed, where each point was representative of a single patient’s measurement for that cell type.

The statistical significance of each correlation was determined by permutation test.

## Acknowledgements

None.

## Author contributions

A.S.M., A.S.R., P.P.L., and R.C.R. conceived of the study. J.R.L. and P.P.L. conducted the experimental analysis. B.O.J. and A.S.M. performed the computational analysis and wrote the first draft. All authors contributed to paper revisions.

## Data Sharing Plan

All analysis was implemented in Python v3.11 and can be found at https://github.com/meyer-lab/tfac-CoH, release 1.0, along with all the experimental data. All other data needed to evaluate the conclusions in the paper are present in the paper or the Supplementary Materials.

