## Supplemental Information for "Systems profiling reveals recurrently dysregulated cytokine signaling responses in ER+ breast cancer patients’ blood"

### Supplement

**Table S1.** Breast cancer patient characteristics and demographics.

| Age | ER | PR | Her2 | Ki67 (%) | TMR Grade | T | N | Stage | Note |
|---|---|---|---|---|---|---|---|---|---|
| 48 | >95 | 40 | - | 1 | 2 | 1b | 0 | Ia | |
| 71 | 95 | 60 | - | 5-10 | 1 | 1B | 1a | IIa | |
| 55 | 90 | 50 | - | 10 | 2 | 1c | 1(m) | Ib | |
| 48 | 90 | 80 | - | 10 | 2 | 2 | 0 | IIb | Chest wall recurrence 5/2021 BRCA 1/2 - |
| 73 | 100 | 90 | - | 5 | 1 | 3 | 0 | IIb | |
| 41 | 90 | 40 | - | 25 | 2 | 1b | 0 | Ia | |
| 42 | > 90 | 70 | - | 10 | 1 | 1b | 0 | Ia | |
| 51 | 90 | 90 | - | 25 | 2 | 2 | 1a | IIb | |
| 35 | >90 | 80 | - | 20-30 | 2 | U | U | U | |
| 65 | >95 | >90 | - | 10-15 | 2 | T2 | 0 | IIa | |

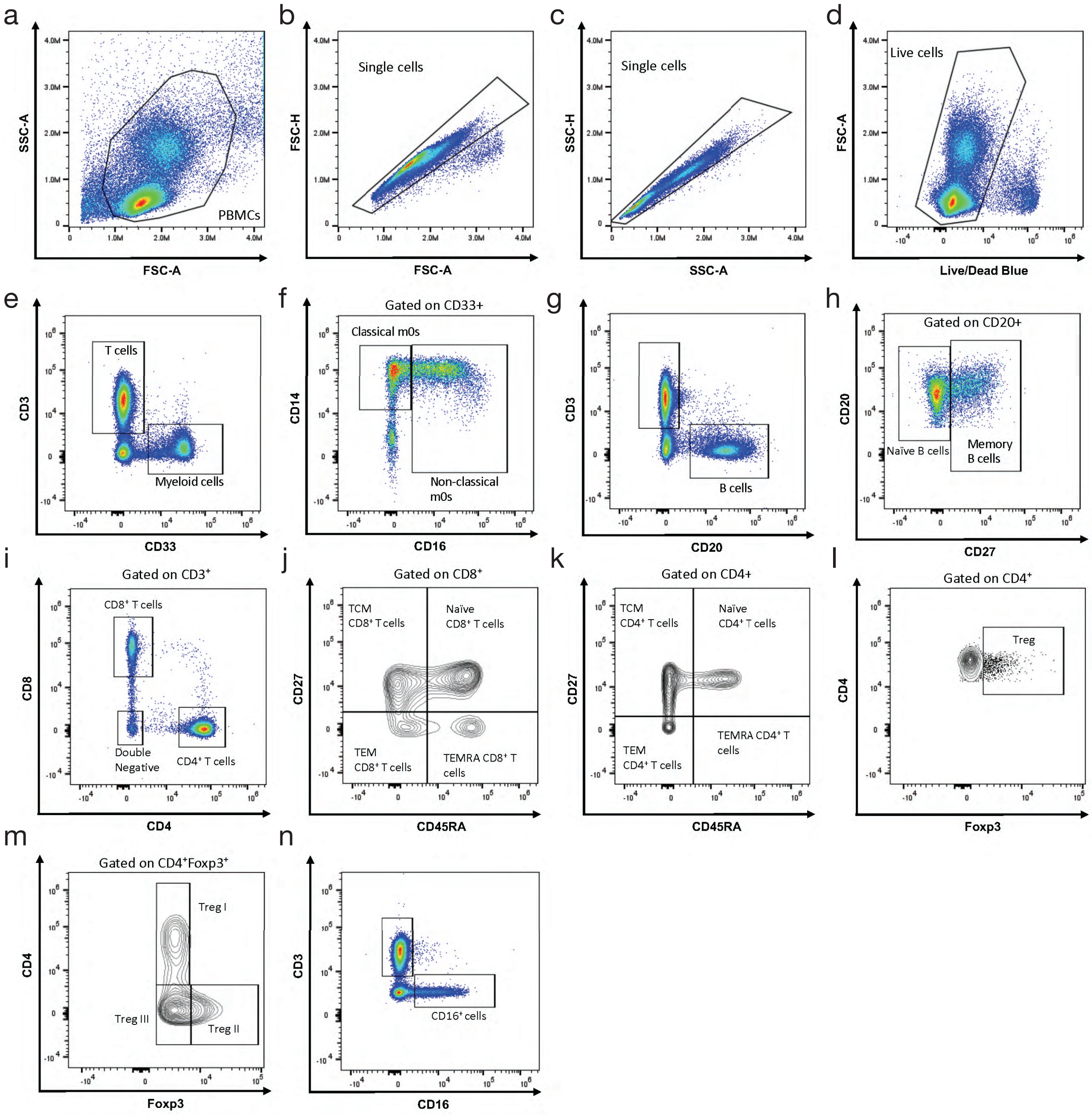


**Fig. S1. Gating strategy for the signaling response and receptor quantification data. (a–d)** Gating for live, single cells. **(e)** Myeloid and T cell gating. **(f)** Gating of classical and non-classical monocytes. **(g)** Gating of B cells. **(h)** Gating of memory and naïve B cells. **(i)** Gating of $CD8^+$, $CD4^+$, and $CD4^-CD8^-$ cells. **(j)** Gating of TCM, naïve, TEM, and TEMRA $CD8^+$ cells. **(k)** Gating of TCM, naïve, TEM, and TEMRA $CD4^+$ cells. **(l)** Gating of Treg cells. **(m)** Gating of Treg 1, Treg 2, and Treg 3 populations. **(n)** Gating of $CD16^+$ population.

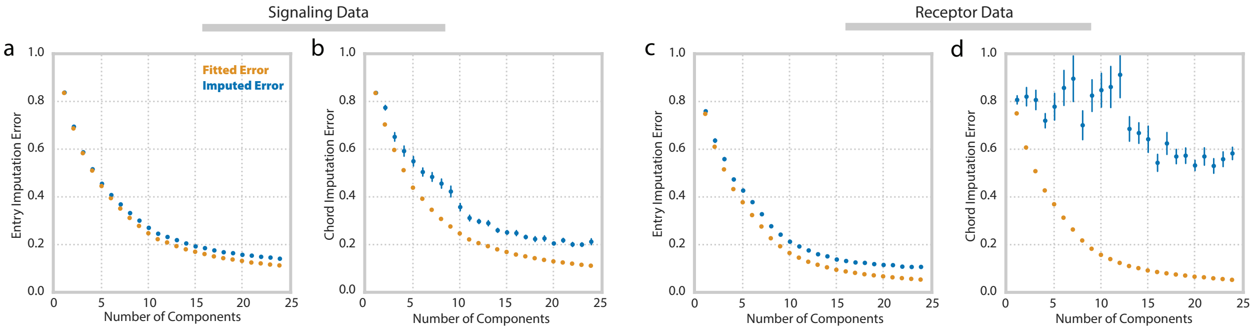


**Fig. S2. CPD can accurately impute the signaling and receptor data. (a,b)** The fitting (gold) and imputation (blue) error using decompositions of varying component numbers for cytokine response dataset. Predictions were made by withholding and subsequently imputing (a) 10% of the entries or (b) 10% of the chords along the patient mode. **(c, d)** The fitting (gold) and imputation (blue) error using decompositions of varying component numbers for the receptor dataset. Predictions were made by withholding and subsequently imputing (c) 10% of entries or (d) 10% of the chords along the patient mode.

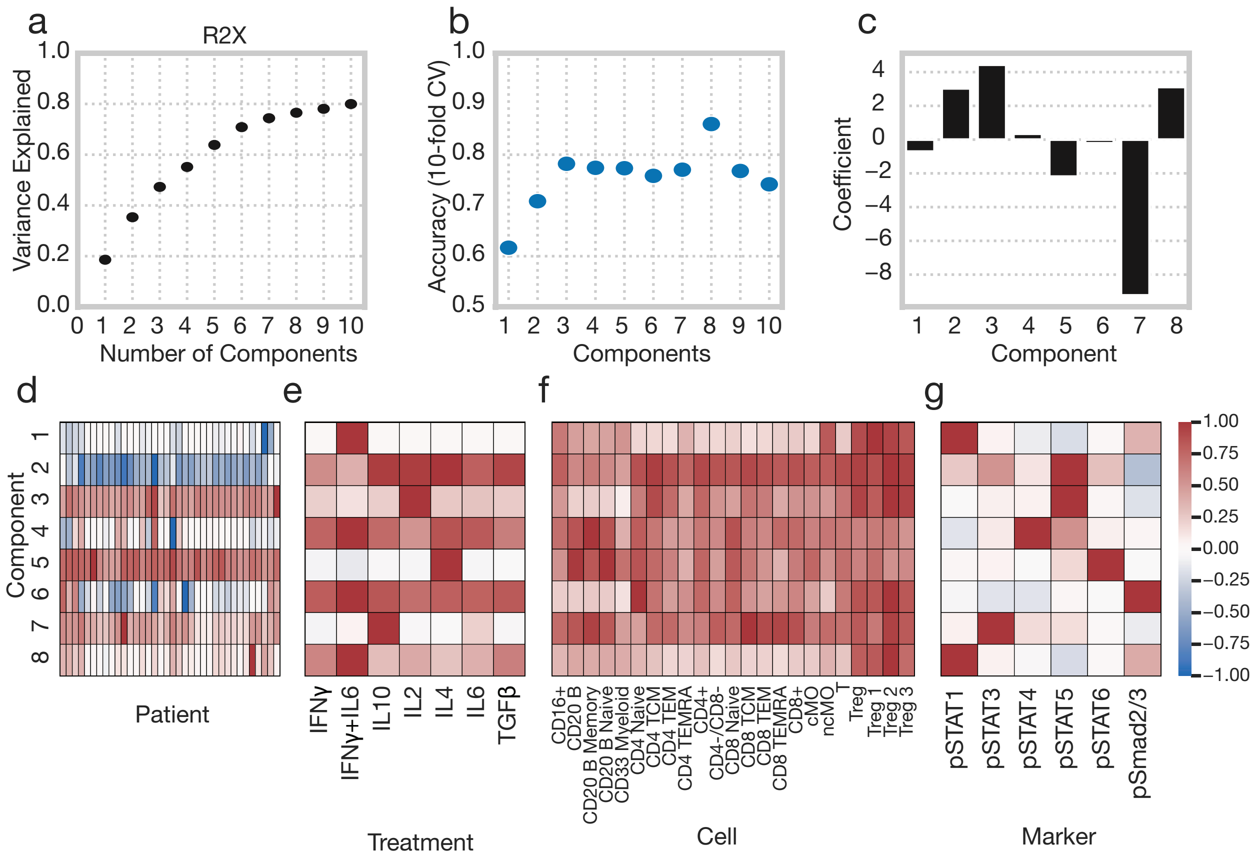


**Fig. S3. Tensor factorization of response data using fold-change metrics reveals similar patterns of dysregulation. (a)** Percent variance reconstructed (R2X) versus the number of components used for decomposition of fold-change signaling dataset. **(b)** Accuracy of logistic regression disease status classification model fit to patient factors for tensor decompositions of varying sizes. Accuracy was determined via 10-fold stratified cross validation. **(c)** Weights of each component for a logistic regression model fit to patient factors using an 8-component decomposition (Healthy = 0, BC = 1). **(d–g)** Component values for each patient (d), treatment (e), cell type (f), and signaling marker (g) collected using a tensor decomposition with 8 components.

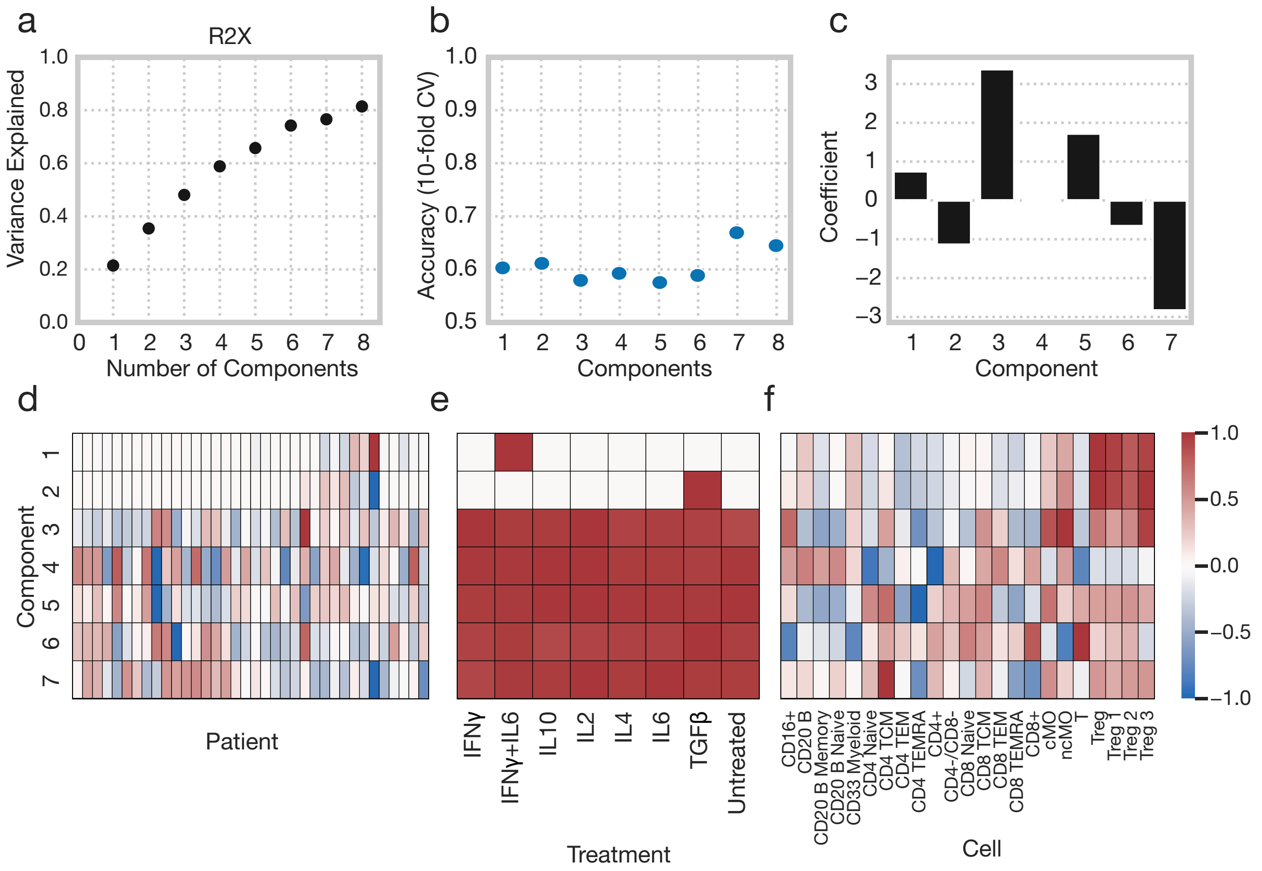


**Fig. S4. Tensor factorization of cell type abundance fails to uncover patterns associated with disease. (a)** Percent variance reconstructed (R2X) versus the number of components used for decomposition of cell type abundance dataset. The abundance of each cell type was calculated by using the percentage of total live lymphocytes which they accounted for. **(b)** Accuracy of logistic regression disease status classification model fit to patient factors for tensor decompositions of varying sizes. Accuracy was determined via 10-fold stratified cross validation. **(c)** Weights of each component for a logistic regression model fit to patient factors using a 7-component decomposition (Healthy = 0, BC = 1). **(d-g)** Component values for each patient (d), treatment (e), and cell type (f) collected using a tensor decomposition with 7 components.

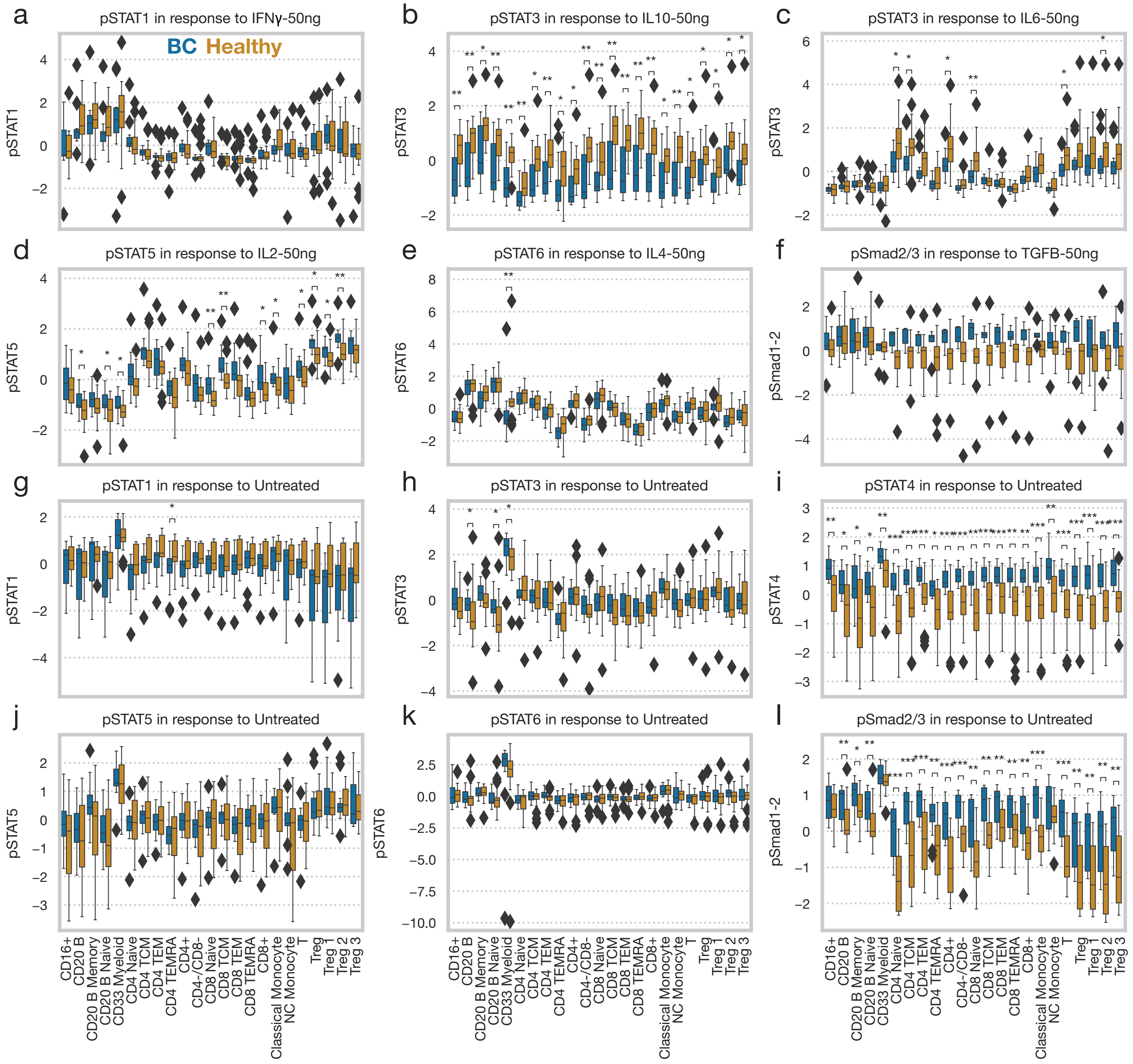


**Fig. S5. Partial panel of signaling responses**. The data was mean centered across all
45 cell types and patients. Statistical significance was determined by two-tailed Mann-Whitney U test. All basal levels are reported as MFI, and induced responses are measured as ΔMFI. *, **, and *** indicate a p-value of less than 0.05, 0.005, or 0.0005, respectively.

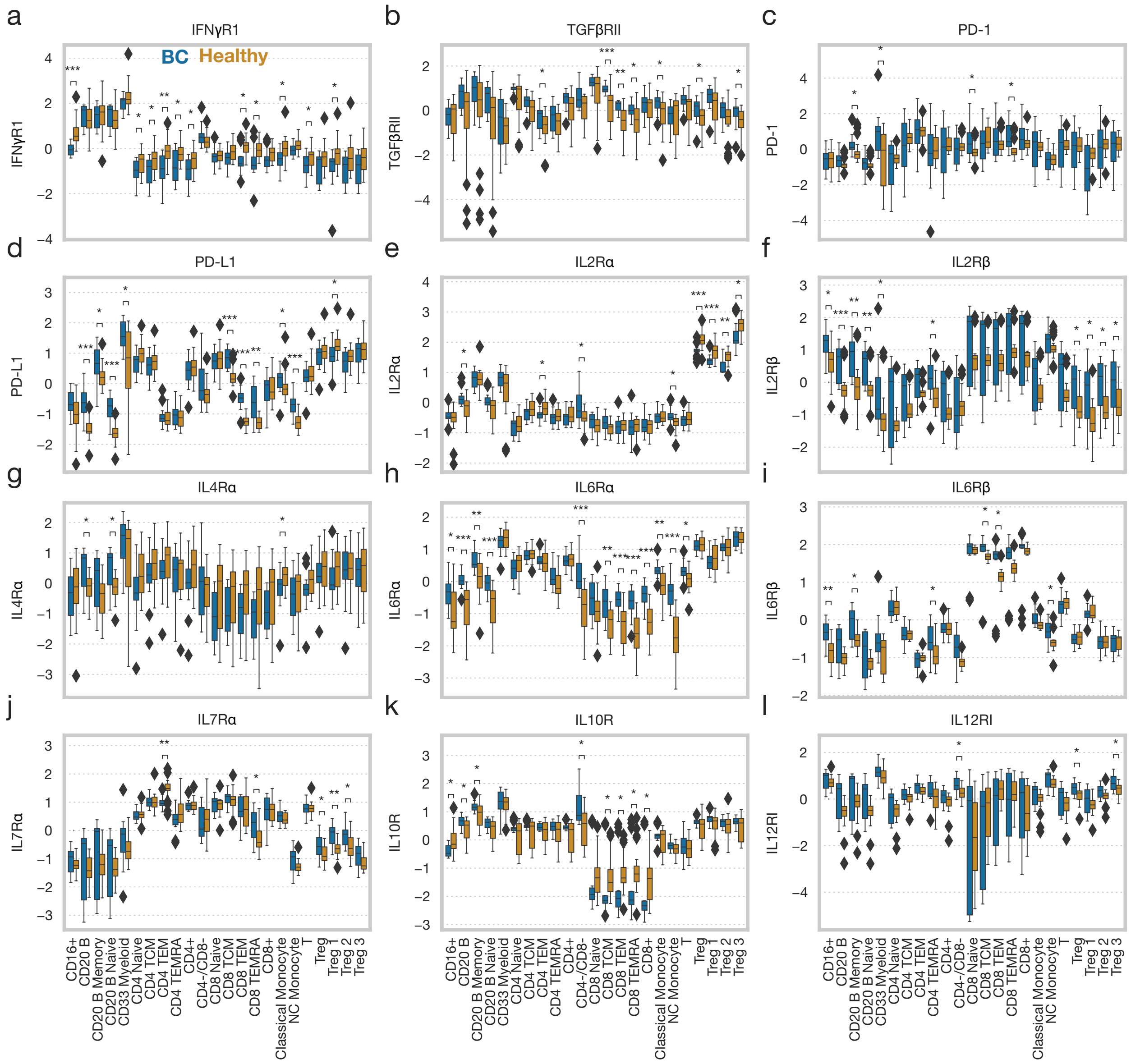


50 **Fig. S6. The full panel of receptor profiling.** The data was mean centered across all cell types and patients. Significance was derived using the Mann-Whitney U test. *, **, and *** indicate a p-value of less than 0.05, 0.005, or 0.0005, respectively.
